# High-content assessment of *Pseudomonas aeruginosa* bacteriophage efficacy reveals host genetic factors involved in phage specificity

**DOI:** 10.1101/2025.04.03.646983

**Authors:** Garrett L. Ellward, Michael J. Bucher, Yuting Zhai, Nathalie S. Munguia, Simone Marini, Rosanna Marsella, Joseph H. Bisesi, Naim Montazeri, Kartikeya Cherabuddi, Kwangcheol C. Jeong, Daniel M. Czyż

## Abstract

*Pseudomonas aeruginosa* infects immunocompromised and hospitalized individuals, resulting in over 500,000 annual deaths. With emerging multidrug resistance and stagnating antibiotic development, alternative antimicrobials are desperately needed. Bacteriophages (phages) offer a promising, effective, and safe alternative. We developed and optimized a high-content liquid assay screen and a stringent assessment of efficacy to isolate and characterize seven novel *P. aeruginosa* phages. Phages were screened individually and in cocktail, inhibiting the growth of over 90% (50/55) of multidrug-resistant clinical strains and ∼75% (102/137) of animal, environmental, and human isolates. When tested in a mouse bacteremia model, the phage cocktail successfully eradicated *P. aeruginosa*. A proteome-wide bi-directional BLAST identified eight proteins that influenced phage infection. The functional analysis of the corresponding genes reveals their putative roles involving genome modification and transcriptional regulation, metabolic processes, and structural components essential for phage docking. Collectively, we have developed a rigorous high-content approach to identify effective phages, which, coupled with functional genomics, revealed genes that affect phage-bacteria interaction.

**Author Summary:** In this study, we explored the potential of bacteriophages (phages) isolated from municipal and hospital wastewater sources for combating multidrug-resistant *Pseudomonas aeruginosa*, an opportunistic pathogen known for posing significant clinical challenges. A rigorous stepwise screen aimed at enhancing specificity against a broad set of 55 clinical *P. aeruginosa* strains allowed us to isolate diverse class phages that can target over 90% of the clinical isolates. Our phage efficacy assessments employed a colorimetric MTT assay to measure the metabolic activity of *P. aeruginosa* strains in response to phage exposure. Notably, the phages demonstrated broad coverage against the *P. aeruginosa* library, with individual phages showing varying degrees of efficacy and a cocktail exhibiting superior inhibitory properties. Further validation using a mouse bacteremia model confirmed the exceptional efficacy of the cocktail, supported by a complete attenuation of clinical signs of infection and a significant reduction of bacterial loads across all organs, supporting their utility as potential phage therapy. Finally, a comprehensive comparative genomic analysis of target bacteria combined with phage efficacy revealed novel genes that are potentially involved in phage infection. These findings provide a foundation for understanding phage-host interactions and pave the way for the development of targeted phage therapies against antibiotic-resistant bacterial infections.

## Introduction

Antimicrobial resistance (AMR) is a growing global issue attributing to millions of deaths each year and crippling the global economy. It is estimated that, by 2050, if no alternatives to current failing therapies are implemented, AMR could be responsible for claiming 10 million lives globally each year and over $1 trillion in medical costs in the United States alone (1,2). The World Health Organization and Centers for Disease Control and Prevention list the ESKAPE pathogens, named for ***E****nterococcus faecium, **S**taphylococcus aureus, **K**lebsiella pneumoniae, **A**cinetobacter baumannii, **P**seudomonas aeruginosa,* and ***E****nterococcus* spp., as priority pathogens for which novel therapies are desperately needed (3,4). *P. aeruginosa*, a gram-negative, motile bacterium found ubiquitously in the environment, typically infects hospitalized patients and individuals with cystic fibrosis, burn wounds, or compromised immune systems (5–7). In 2019, *P. aeruginosa* was one of six pathogens responsible for approximately 73% of the 1.27 million deaths attributed to AMR (8). As one of the more common nosocomial pathogens, often complicating medical treatment, infections by *P. aeruginosa* have reported mortality rates upwards of 40% (9,10). With intrinsic resistance to antimicrobials, rapid evolution, and acquisition of pan-drug resistance*, P. aeruginosa* continues to be one of the most pressing AMR threats, highlighting the dire need for developing new therapeutic strategies (11–13).

Bacteriophages (phages), viruses that can specifically target and kill bacteria, offer a promising alternative or supplement to current antimicrobial therapies. First discovered over 100 years ago, the antimicrobial properties of phages have been extensively studied, and historically, phages have been used clinically to treat infections in countries like Georgia, Poland, and Russia (14,15). Despite their efficacy, phage therapy was abandoned in Western medicine for several reasons, including the lack of standardized studies, the availability and mass production of effective antibiotics, and political strife (16,17). Currently, phage therapy is reemerging as a possible treatment for AMR infections. In the United States, phage therapy has been increasingly more common in the past decade; however, authorization for therapy is still rare and is reserved for compassionate use cases after all other options have been exhausted (18,19). Despite preliminary successes in clinical applications, the acceptance of phage therapy as a standard treatment remains hindered by the lack of clinical trials and pre-clinical animal models.

Given the extensive resistance of *P. aeruginosa* to current antimicrobial treatments, the high rate of recurrent infections (20), and not being a commensal resident of the human microbiota, this pathogen has become an attractive target for phage therapy. Several case studies have already been conducted utilizing *P. aeruginosa* phages (21–25); however, not all were successful, primarily due to the emergence of resistance and sub-optimal efficacy of phage combination (26). Despite encountering challenges (27), phage therapy remains a prospective option for AMR treatment.

Several important aspects of phage-host interaction are critical for successful phage therapy. Namely, the host specificity influenced by phage receptors at the bacterial membrane (28), the capacity of phages to lyse host cells facilitated by the mechanisms favoring the lytic infection cycle (29), the role of superinfection exclusion (30), and bacterial factors involved in phage replication, defense, and release (31). Additionally, factors that drive phage-host co-evolution are likely to drive the adaptability and specificity of phages in therapeutic applications. Therefore, understanding host factors that influence phage infection is crucial to improving phage utility as an antimicrobial tool and predicting phage efficacy.

Traditionally, the killing potential of phages is assessed via plating against bacterial hosts on solid media (32). While a proven method to test phage efficacy, utilization of solid media assays can be relatively slow and limited in scope due to time and material constraints. Additionally, a solid medium may not recapitulate phage dynamics *in vivo*. For this reason, a need for a high-content pipeline of phage isolation and characterization has emerged to streamline the preparation of phage for therapeutical applications. Previous work has built the foundations of using colorimetric assays and optical density measurements to test phage efficacy (22,33,34). In this study, we developed a stringent phage infectivity assessment method using a tetrazolium-based colorimetric assay (liquid assay) in conjunction with a high-throughput OmniLog® system and a combination of three metrics: Reduction in metabolic activity (RiMA), time until relative units (TURU), and area under the curve (AUC). We have isolated seven novel *P. aeruginosa* phages and employed our assessment method for their characterization using a comprehensive collection of 184 human, animal, and environmental isolates. We confirmed the efficacy of the most effective phage combination using a mouse model, completely rescuing animals from lethal *P. aeruginosa* bacteremia, resulting in no clinical signs of infection. Employing comparative genomics in combination with our phenotype-genotype data, we analyzed whole-genome sequences of 132 *P. aeruginosa* strains for genes involved in phage infection. A whole-genome bi-directional BLAST analysis yielded a total of 19 genes; knockout mutations for five of these genes confirmed an increase in phage sensitivity, and a knockout of four increased phage resistance. Our study adopts a high-content pipeline for the isolation and identification of phages with strong clinical potential and yields host genes that affect phage infection.

## Results

### Phage Isolation and Characterization

We collected untreated raw wastewater from several locations for phage isolation. Primarily, water samples were collected from the UF Water Reclamation Facility. To increase the range of the isolated phages and likely their specificity against clinical isolates, we tapped into the UFHealth Shands Hospital wastewater system. Using these two sources, we isolated seven novel phages. Our initial standard three-round isolation yielded a heterogeneous population of phages, as revealed by transmission electron microscopy (TEM) imaging. As such, we increased the standard number of phage isolations from three to nine rounds before beginning to characterize the phages using liquid assay (Fig 1A). We took an iterative screening approach to isolate phages effective against a collection of 55 clinical isolates of multidrug-resistant (MDR) *P. aeruginosa* (S1 Table). We screened the first isolated phage against all the 55 strains, selected the next *P. aeruginosa* strain that is resistant to the isolated phage, and used it as bait to isolate another phage. This stepwise isolation method was continued until ∼90% of the strains in the bacterial collection were targeted by the isolated phages. Using this approach, we have isolated seven phages: Pa001_Violet, Pa002_Juniper, Pa003_Lilly, Pa004_Willow, Pa005_Rush, Pa006_Rose, and Pa007_Sage. Next, we employed a combination of TEM and genomic analysis to determine the morphology of each phage, revealing a diverse array of classes, including phiKZ-, Bruynoghe-, Pakpuna-, Lituna-, Septimatre-, and Citex-like viruses (Fig 1B). The genome sequencing of each phage confirmed their closest relatives and taxonomic classification (Fig 1C, Table 1, S2 Fig). A comprehensive analysis of each genome revealed a single phage, Pa006_Rose, carrying a predicted integrase and a Zeta toxin (Table 1).

**Fig 1.**
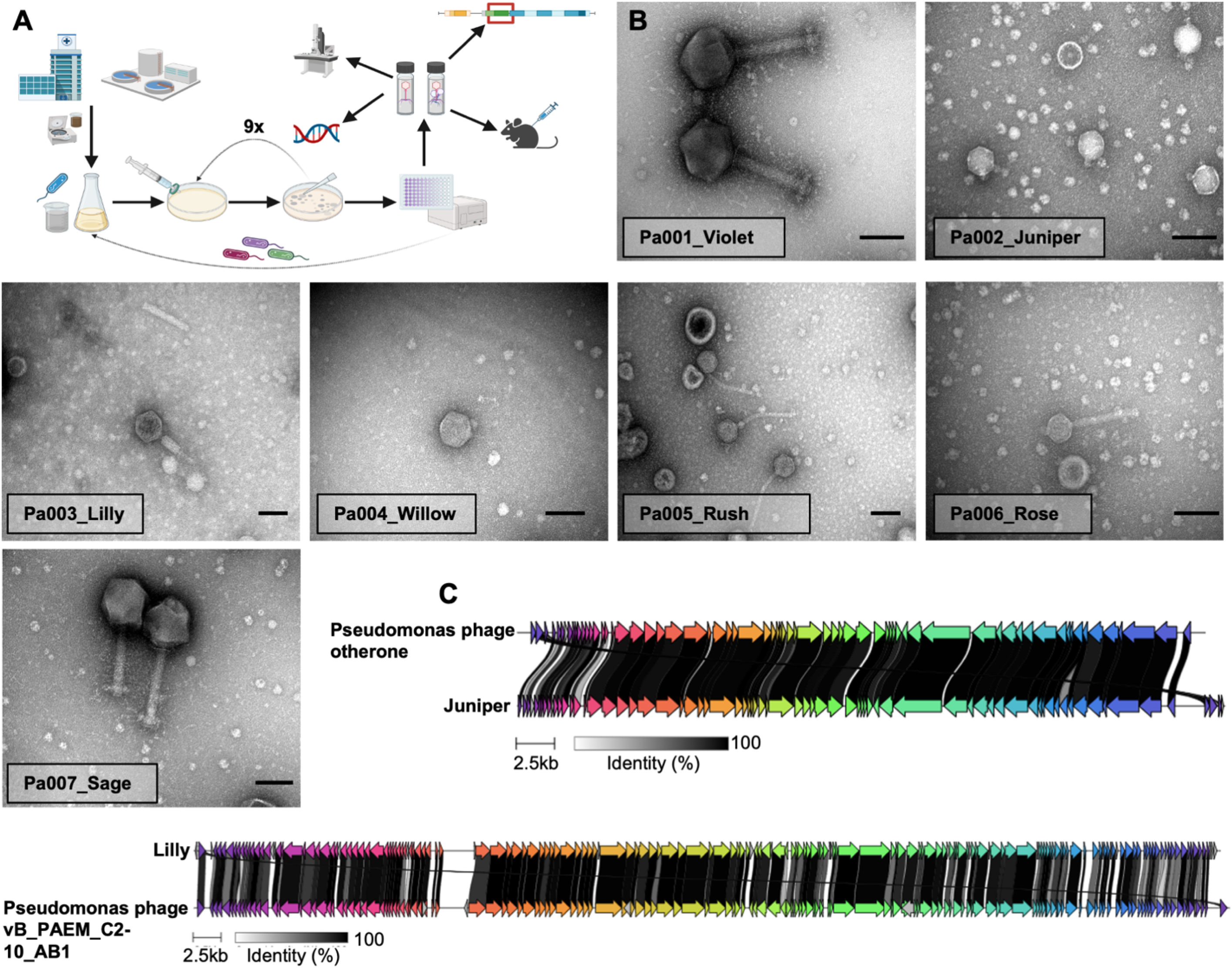
Isolation and characterization of novel phages. A) Graphical overview of the experiment. Seven novel phages were isolated from municipal and hospital wastewater in a stepwise manner. After initial screening against a clinically relevant bacterial collection, phages were combined equally into a cocktail to test against our comprehensive collection using the liquid assay. Individual phages were further characterized using TEM and Illumina sequencing. After characterization, the cocktail was used to treat mice *in vivo,* and bacterial genes involved in phage infection were identified and confirmed using a bi-directional BLAST and confirmed using a *P. aeruginosa* transposon mutant library. B) TEM images of seven phages isolated from wastewater. Phages were imaged using an FEI Tecnai Spirit 120 kV TEM. All scale bars represent 100 nm. C) Clinker alignment of Pa002_Juniper against its closest relative. Sequencing and imaging confirmed homogenous phage stocks.

**Table 1.**
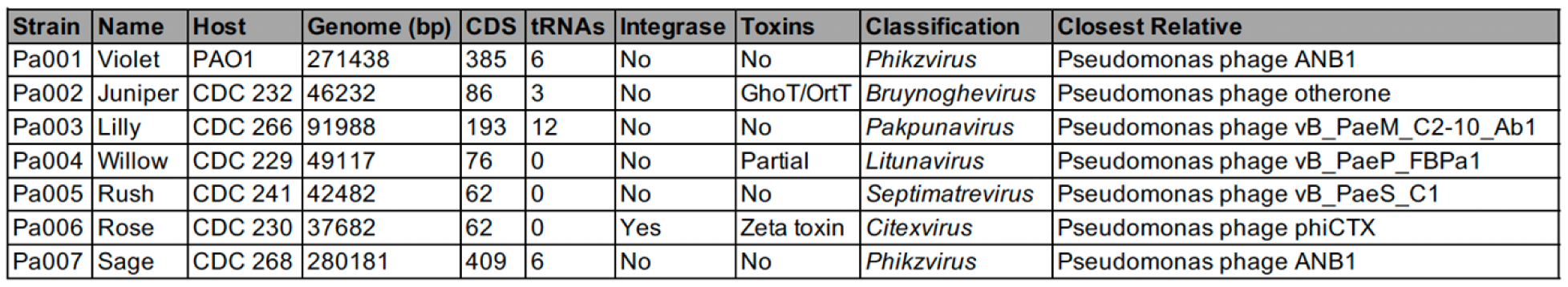
Phage genomic information and closest relatives.

### Individual Phage Efficacy

Upon genotypic and phenotypic characterization of pure phages, we began screening each of the seven isolates in a high-throughput *in vitro* (3-(4,5- Dimethylthiazol-2-yl)-2,5-Diphenyltetrazolium Bromide) (MTT) colorimetric assay to determine their efficacy against *P. aeruginosa*. MTT, when reduced during bacterial metabolic respiration, turns from yellow to purple, a color change that is detected by the OmniLog® reader to gauge the relative metabolic activity of bacteria in 96-well plates (S3 Movie). Although we used a specialized OmniLog® reader, any spectrophotometer can be used for smaller-throughput applications. The *P. aeruginosa* library contained 55 MDR, including extensive and pandrug-resistant clinical isolates of *P. aeruginosa* obtained from the CDC’s AR Isolate Bank (Panel ID 12). Additionally, we included *P. aeruginosa* PAO1 and mPAO1 for a total of 57 bacterial isolates that were screened with each phage. Sensitivity and resistance to phage were determined by assessing significance across three separate metrics: AUC, RiMA, and TURU (Fig 2). The AUC values were calculated using an approximation of the Riemann’s sum. The total area under the curve generated during the first 12 hours of growth was analyzed for any significant difference between no phage and phage groups (Fig 2A). Since MTT is reduced during bacterial metabolic processes, the change in color is a representation of the metabolic activity of the bacterium and, hence, its viability. The RiMA score represents the difference between the metabolic rates of the no-phage and phage groups (Fig 2A, Fig 2D). To be considered significant, we used a cutoff score equal to or greater than ten, which is about three standard errors from the average PAO1 control (S4 Fig). This RiMA score cutoff was chosen based on our initial screens, revealing an average error in the metabolic activity in no phage groups. The last metric, TURU, examines the time it takes for the signal to reach a certain threshold during the experiment. Specifically, how long it takes before the metabolic rate reaches 100, 200, 300, and 400 Relative Units (the output of the OmniLog® system) (Fig 2A). Though RiMA and AUC analyses examine the infection kinetics during the first 12 hours, TURU analysis was extended to 22 hours, as some bacteria require additional time to reach 400 Relative Units. An example of phage-sensitive and resistant bacterial isolates is represented in Fig 2A. A strain was deemed sensitive to a phage if all three metrics significantly differed from a no-phage control. This triple-metric approach ensures stringent criteria and sensitivity for phage selection. To ensure that our assessment reflects phage killing efficacy, we cultured *P. aeruginosa* PAO1 at different multiplicity of infection (MOI) of the Pa007_Sage phage over 22h, followed by serial dilutions and plating for colony-forming units (CFU) (S5 Fig). The results revealed that despite the similar end-point reduction of MTT, the shift in RiMA resulted in the nearly complete eradication of *P. aeruginosa*. These results support our approach and the choice of the three-metric assessment.

**Fig 2.**
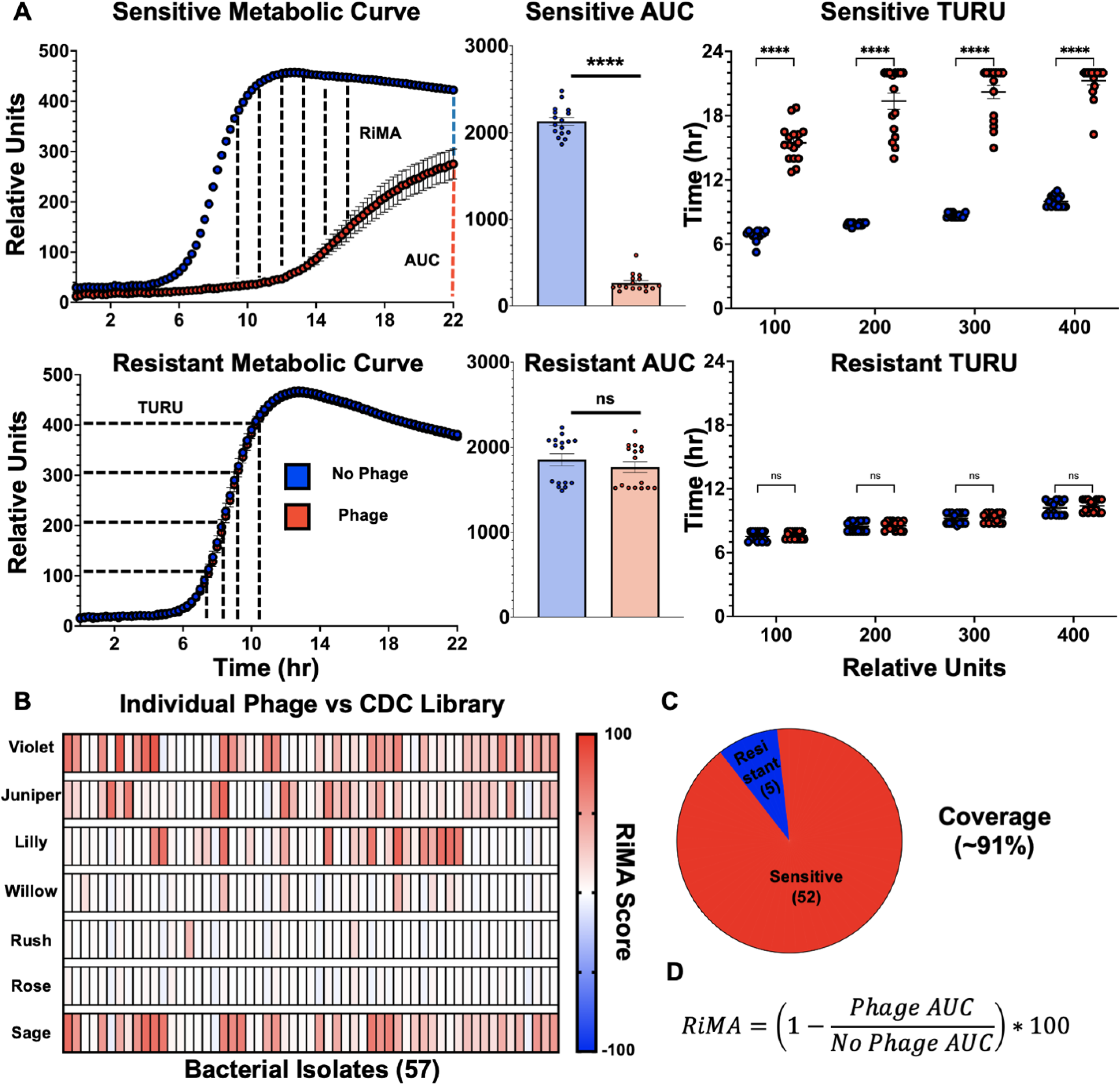
Initial bacteriophage screen against *P. aeruginosa* CDC isolates. A) Examples of sensitive and resistant bacterial isolates after 12 hours of exposure to phage. Sensitivity or resistance is determined by examining significance across three metrics: RiMA, AUC, and TURU analyses. B) Heatmap representation of each phage against the initial CDC library. In total, 57 different bacterial isolates were tested against all seven phages. Red indicates more killing, and blue indicates less. C) Library coverage using individual phage results. D) The formula for calculating RiMA. Error bars represent standard error of the mean (n=16). Statistics were performed using a Student’s *t*-test (AUC) and Multiple *t*-test (TURU). (*, p<0.5; **, p<0.1; ***, p<0.001; ****, p<0.0001; ns, not significant).

Four of the seven phages isolated—Violet, Juniper, Lilly, and Sage—showed strong inhibition towards *P. aeruginosa*, targeting 32, 28, 22, and 33 different isolates, respectively (Fig 2B, Table 2). The remaining phages, though not as virulent across all strains, were able to target bacteria that other phages could not, collectively increasing the overall coverage of the CDC library. Only five bacterial isolates were resistant to every phage (Fig 2C). While each individual phage targeted anywhere between 7 to almost 58% of isolates, collectively, they targeted over 90% of the *P. aeruginosa* isolates, demonstrating a robust specificity against clinically important MDR strains (Fig 2C).

**Table 2.**
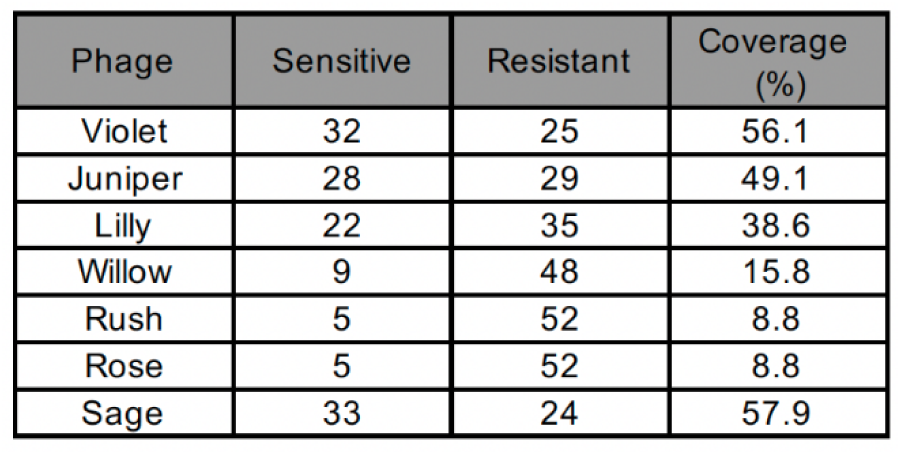
Phage infection summary against CDC bacterial isolates.

| Phage | Sensitive | Resistant | Coverage (%) |
| --- | --- | --- | --- |
| Violet | 32 | 25 | 56.1 |
| Juniper | 28 | 29 | 49.1 |
| Lilly | 22 | 35 | 38.6 |
| Willow | 9 | 48 | 15.8 |
| Rush | 5 | 52 | 8.8 |
| Rose | 5 | 52 | 8.8 |
| Sage | 33 | 24 | 57.9 |

Our colorimetric *in vitro* screening method measures metabolic activity, which generally serves as a reliable indicator of bacterial viability, though it does not directly assess it. To confirm that RiMA corresponds to the loss in bacterial viability and reflects phage lytic properties, we combined all phages equally in cocktail and tested its efficacy using the described liquid assay compared to spotting assay. The combination of phages was spotted at 1×10^6^ plaque forming units (PFU) onto a 0.7% top agar layer on tryptic soy agar (TSA) plates coated with each of the 57 *P. aeruginosa* hosts (S6 Fig).

After approximately 8-12 hours of incubation, the plates were examined for plaque formation. Bacterial responses to the cocktail were classified as sensitive (clear plaques), intermediate (some plaques), or resistant (no plaques). While the spotting assay confirmed the sensitivity of 46 of the 57 strains, six strains that were resistant in liquid tested either sensitive or intermediate on solid medium, and five became resistant when tested on solid. While these results validate the MTT colorimetric assay as a viable approach to assess phage efficacy, they also reveal the differences in phage specificity influenced by the growth medium (liquid vs solid) of the bacterial host.

### Cocktail Efficacy

We assessed the efficacy of the cocktail against a broad spectrum of *P. aeruginosa* isolates and determined host-range specificity by testing the combination of our seven phages. To maintain the same MOI, we prepared a final cocktail concentration of 1×10^4^ PFU/mL, which is the concentration equivalent to that used for the assessment of each individual phage. We used an expanded collection of 137 animal and human isolates of *P. aeruginosa* and an additional 47 non-*aeruginosa* species (S7 Table). The phage cocktail showed an exceptional specificity to *P. aeruginosa* strains, as none of the non-*aeruginosa* species were targeted, which was evident from our three stringent metrics (S7 Table, S8 Fig).

The phage cocktail showed a robust efficacy against *P. aeruginosa* (Fig 3A). Average RiMA scores against the two controls, PAO1 and mPAO1, and 55 CDC *P. aeruginosa* strains increased upon treatment with the phage cocktail compared to individual phages, as indicated by the darker orange cells (more phage activity). The liquid screening revealed the specificity of the cocktail against approximately 75% (102/137) *P. aeruginosa* isolates (Fig 3B). The cocktail strongly inhibited clinical human and animal isolates (Table 3). Despite each individual phage being 1/7th of its titer used for individual phage testing, the combination of the seven phages improved anti-*P. aeruginosa* efficacy. In fact, many isolates from the CDC *P. aeruginosa* collection showed higher RiMA scores when treated with the cocktail compared to individual phage (Fig 3C), suggesting complex interactions such as cooperative enhancement, suppression of resistance mechanisms, or other synergistic properties (35). Collectively, both individual phages and the cocktail showed exceptional specificity and efficacy against *P. aeruginosa* isolates when tested *in vitro*.

**Fig 3.**
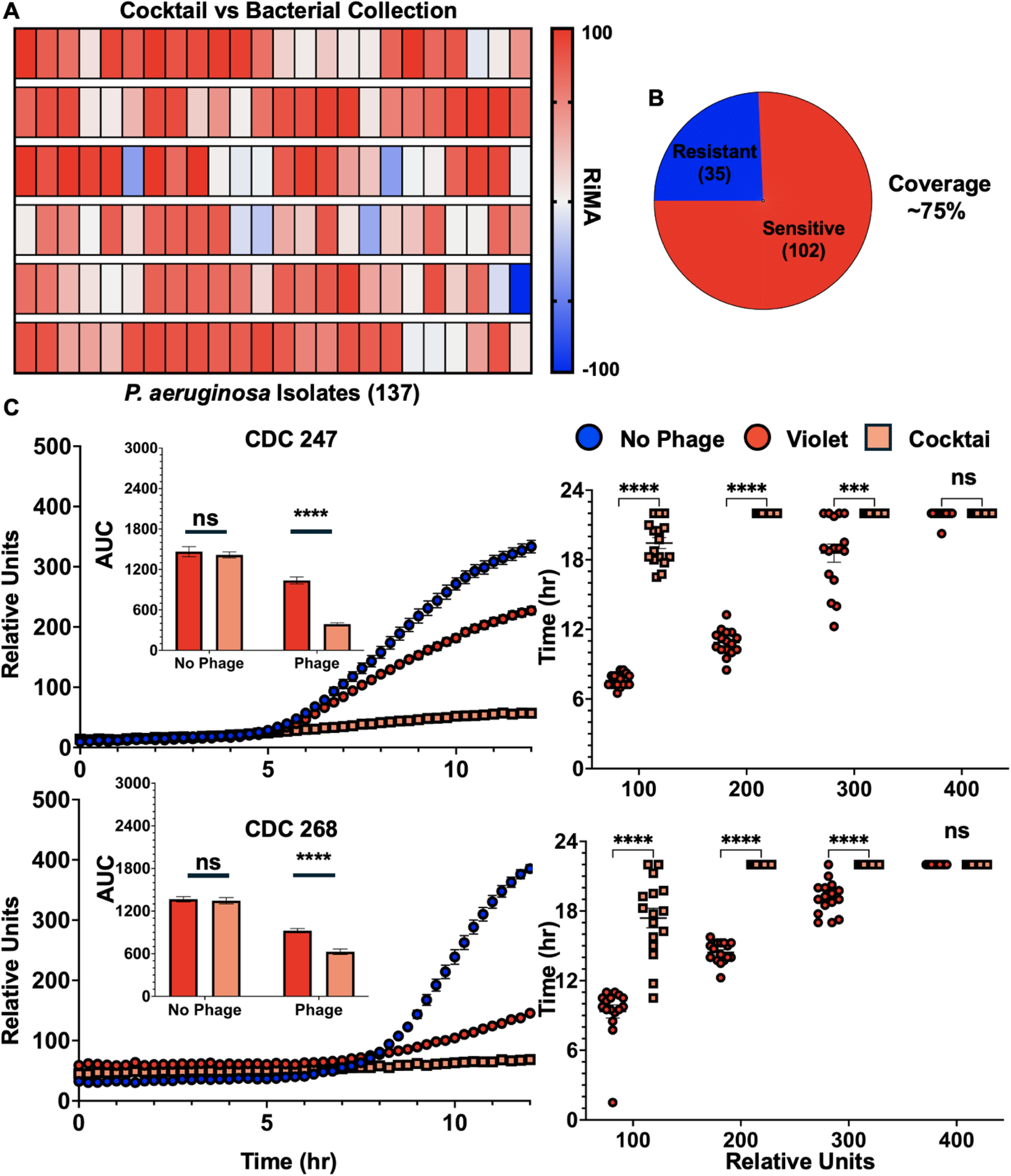
Cocktail efficacy against 137 unique *P. aeruginosa* isolates. A) Calculated RiMA scores for the bacteriophage cocktail against the entire strain collection. Red cells indicate more phage activity, while blue indicate less. A total of 137 unique *P. aeruginosa* isolates were tested. B) Summary of phage cocktail coverage. C) CDC isolates showed higher RiMA scores when treated with the cocktail compared to individual phage treatment. Error bars represent standard error of the mean (n=12). Statistics were performed using a Student’s *t*-test (AUC) and multiple t-test (TURU). (*, p<0.5; **, p<0.1; ***, p<0.001; ****, p<0.0001; ns, not significant).

**Table 3.**
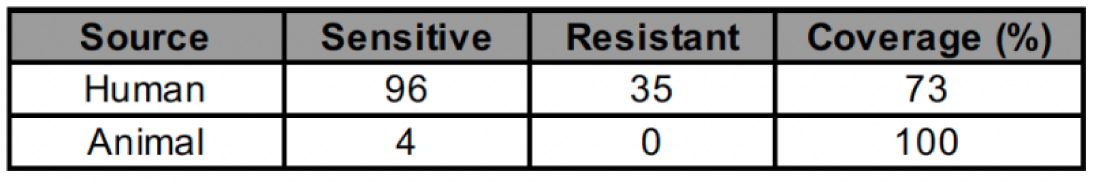
Phage efficacy by isolation source.

### Mouse Model

Following promising *in vitro* results, we aimed to test the efficacy of the experimentally derived phage cocktail in an *in vivo* mouse peritonitis model. Ten 8-12-week-old female C57BL/6 mice were selected to test cocktail efficacy (Fig 4). At time zero, all mice were inoculated with 1×10^6^ CFU of *P. aeruginosa* PAO1 via intraperitoneal (IP) injection. Thirty minutes post-infection, after allowing the bacteria to disperse throughout the body, the treatment group received IP injections of 1×10^9^ PFU of the phage cocktail consisting of all but Pa006_Rose due to the presence of a putative integrase, while the control group was injected with sterile phage buffer. The phage cocktail was re-administered twice every 12 hours following the initial bacterial inoculation (Fig 4).

**Fig 4.**
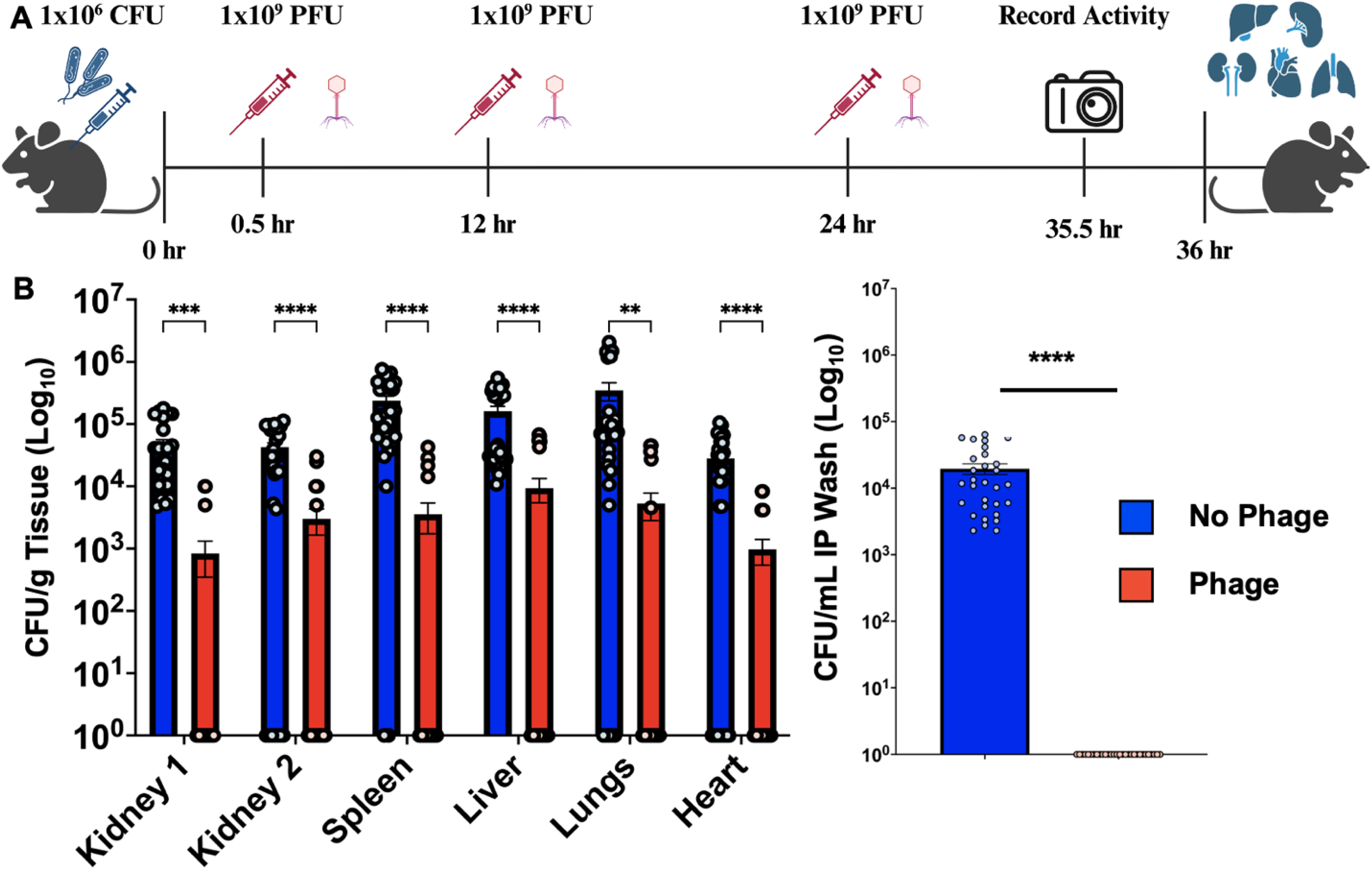
Murine model using cocktail. A) Experimental overview. At time zero, mice were inoculated intraperitoneally with 1×10^6^ CFU *P. aeruginosa* PAO1. Thirty minutes later, phage mice were injected with 1×10^9^ PFU phage cocktail. Reapplication of treatment occurred every 12 hours thereafter. Mice were sacrificed at 36 hours. Figure cartoon created with Biorender.com. B) Bacterial load enumeration in CFU/gram for organs or CFU/mL for intraperitoneal (IP) wash. Error bars represent standard error of the mean (n=30). Statistics were performed using a multiple t-test (*, p<0.5; **, p<0.1; ***, p<0.001; ****, p<0.0001) (ns, not significant).

After 36 hours, the mice were euthanized and dissected to extract kidneys, liver, spleen, lungs, and heart for bacterial load assessment. Additionally, the IP cavity was washed with phosphate-buffered saline (PBS) for bacterial enumeration. Phage treatment resulted in a significant decrease in bacterial load across all organs. Nearly every single phage-treated mouse had completely cleared the *P. aeruginosa* infection, as no colonies were recovered from most phage-treated mice. However, a single mouse in the phage-treated group had detectable CFU counts, possibly due to the initial phage injection being inoculated into the intestine, localizing the phage treatment and preventing the killing of bacteria. Nonetheless, after 36 hours, the total bacterial content across organs for this single mouse was significantly lower than buffer-injected mice, with p ≤ 0.01 (S9 Fig). Immediately before mice were sacrificed, a video was recorded to capture the striking difference in physical activity and clinical signs of infection between the control and phage treatment groups (S10 Movie). While all control mice exhibited severe clinical signs of infection manifested by reduced activity, ruffled fur, and hunched posture, all phage-treated mice had no signs of infection, including the single mouse that had detectable CFU counts (S9 Fig). Our *in vivo* results confirm the exceptional performance of the six-phage formulation in the treatment of bacteremia.

### Identification of Genes Affecting Phage Infection

To identify bacterial genes involved in phage infection, we leveraged our extensive phenotypic and genotypic data. First, using 500 single-copy conserved genes, we constructed a phylogenetic tree from 125 good-quality whole genome sequences of *P. aeruginosa* (Fig 5). We superimposed our phenotypic data onto the phylogenetic tree based on 102 phage-sensitive and 35 phage-resistant isolates of *P. aeruginosa* (Fig 3). Interestingly, a single clade emerged, containing a cluster of seven *P. aeruginosa* strains that exhibited resistance to the phage cocktail (Fig 5). We employed these seven resistant strains, along with three sensitive strains, PAO1, CDC 264, and K016, of which the two latter cluster with the seven resistant strains. We ran a whole proteome bi-directional BLASTp analysis on these ten isolates, looking for proteins with sequence similarity scores of 50% or less between the resistant and sensitive strains. We selected protein candidates that were below the threshold across all bacteria in the resistant and sensitive clusters. In total, we identified 19 proteins with low sequence similarity scores between phage-resistant and sensitive strains (Table 4). The low sequence similarity of these proteins suggests their potential role in phage infectivity. To ensure that our cutoff of 50% did not lead to the selection of proteins other than the identified candidates, we aligned all proteins across the seven resistant and three sensitive *P. aeruginosa* strains. As expected, the sequences of the sensitive strains are different from the resistant strains and cluster together (S11 Fig).

**Fig 5.**
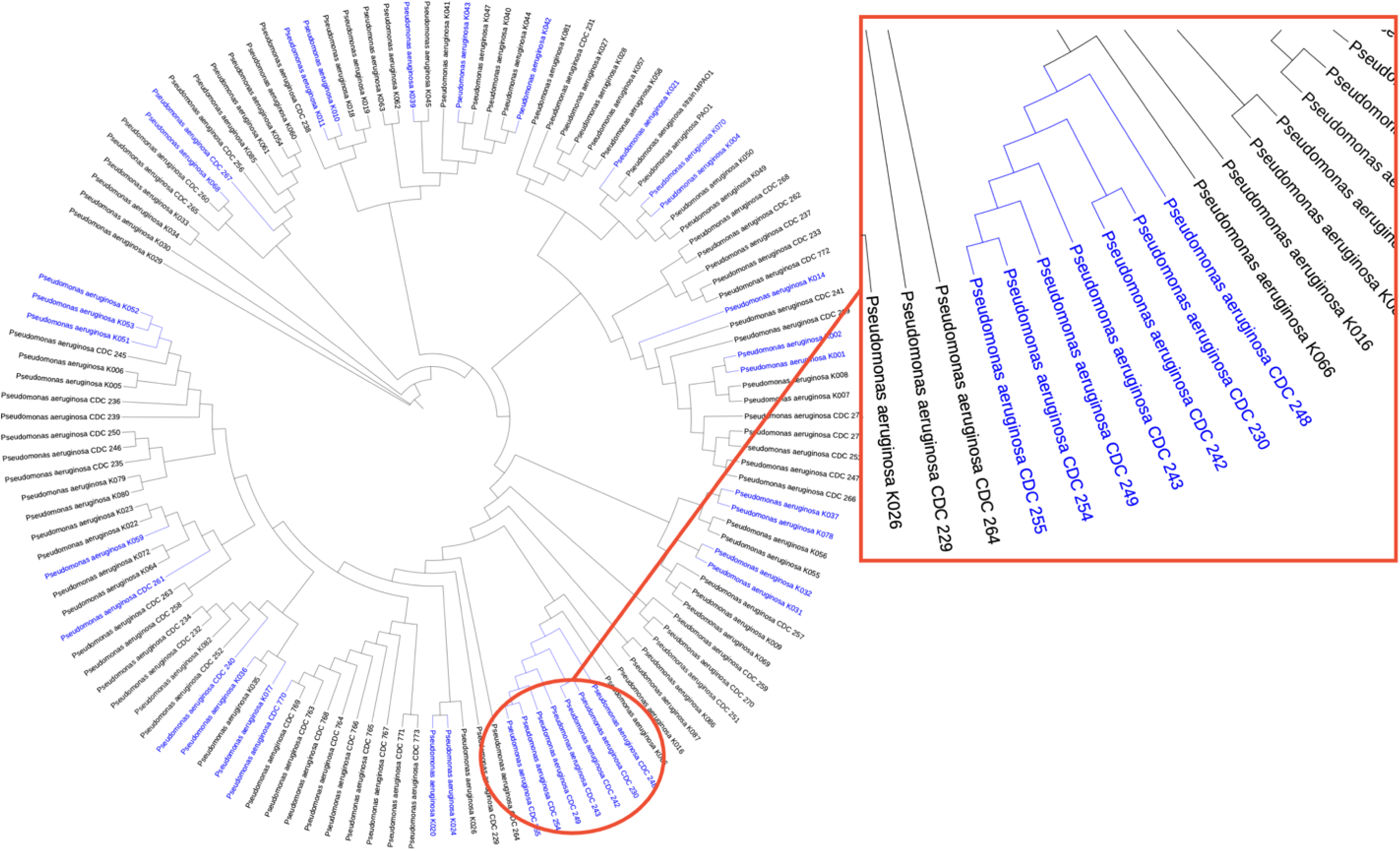
Phylogenetic analysis of *P. aeruginosa* isolates. Using 500 single-copy genes, a phylogenetic tree of 125 *P. aeruginosa* isolates was constructed. A clade of seven resistant CDC isolates was discovered and analyzed alongside two sensitive isolates via a bi-directional BLASTp. Names in blue highlight resistant strains.

**Table 4.**
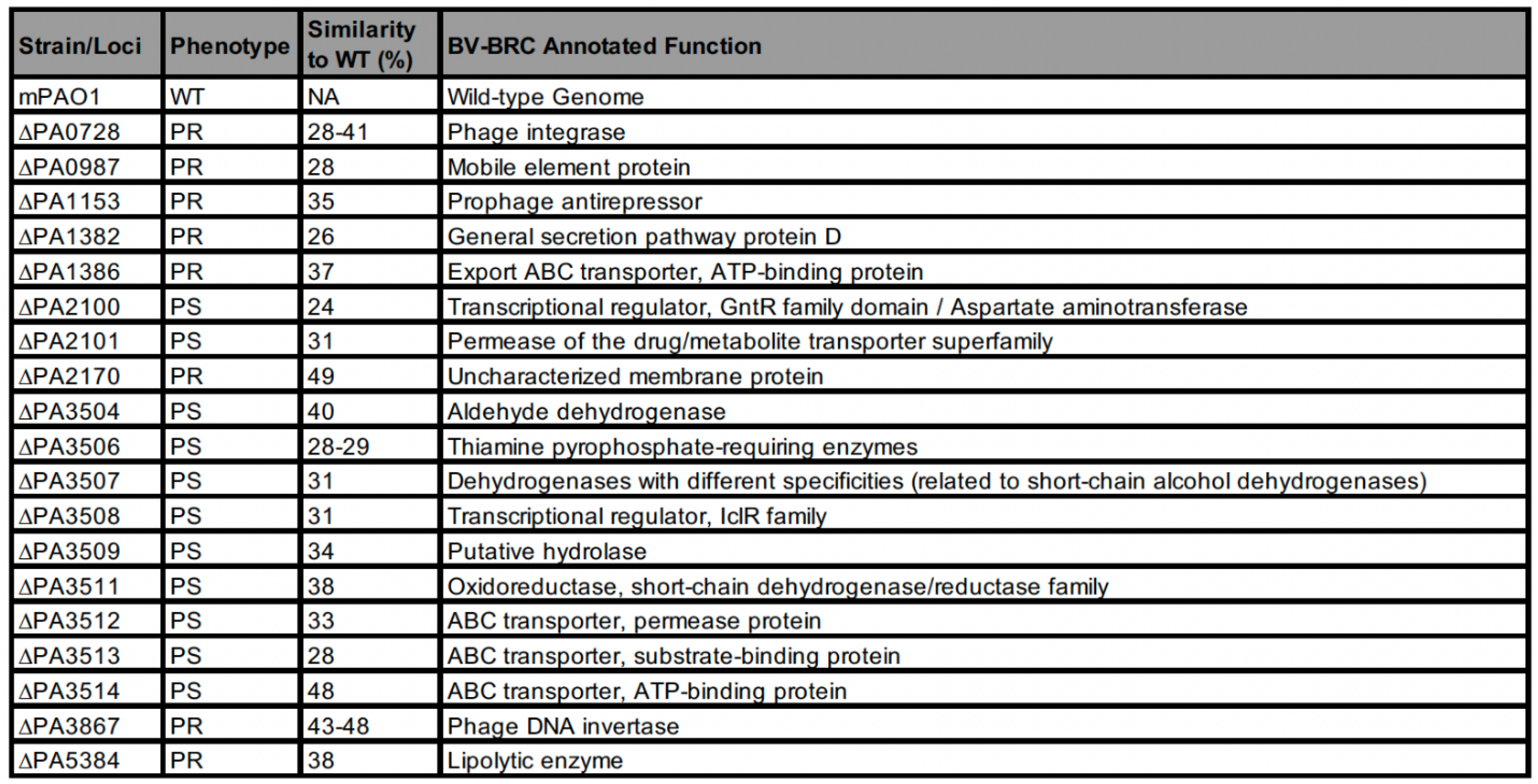
Putative *P. aeruginosa* genes involved in phage infection. Genes with less than 50% similarity to the wild type PAO1 were identified from the bi-directional BLASTp. “PS” denotes strains that carry a mutation in genes resulting in a phage-sensitive strain; “PR” denotes phage-resistant strain.

To confirm the role of these bacterial components in phage infection, we used *P. aeruginosa* transposon knockout mutants for each of the genes coding for the identified proteins. We employed the liquid assay to test the specificity of the two most potent phages, Pa001_Violet and Pa007_Sage, against mutant alleles (two for most genes) for each of the 19 target genes using an initial MOI of 0.5. The interactions between mutant strains and phages were assessed using three metrics: RiMA, AUC, and TURU. If a mutant exhibited a significant difference in phage specificity based on the three requirements of the metrics relative to the control wild-type, it was deemed as a putative bacterial component that affects phage infection. In total, mutants of eight genes exhibited a significant difference in their interaction with the phages, highlighting their potential role in phage infection (Fig 6A, S12 Table). Of these eight genes that exhibited significant differences in phage infection, five (PA1382, PA3507, PA3508, PA3512, PA3867) significantly increased sensitivity towards Violet, Sage, or both phages; similarly, four genes (PA0728, PA0987, PA2100, and PA3867) significantly increased resistance towards the selected phages (Fig 6B). Interestingly, several genes (PA3504 through PA3514) identified by the bi-directional BLAST are located within the same operon. Liquid assay of their corresponding mutants revealed three genes (PA3507, PA3508, and PA3512) that exhibited enhanced sensitivity to phages (Fig 6C). The close proximity of these genes suggests common regulatory mechanisms, further supporting their potential role in modulating phage infection. To better understand potential interactions between the identified genes and phage infection, we summarized the cellular localization and predicted functions of the candidate genes (Fig 6D). We ran a bacterial localization prediction tool (PSORTb v3.0.3) to determine the subcellular localization of each of the predicted gene products (36) (Fig S13 Table). Products of five of the eight predicted genes are located within the cytoplasm but have predicted functions that can affect interaction with phages, such as transcriptional regulators, proteins involved in fatty acid synthesis, and possible prophage genes, which could be involved in superinfection exclusion. PA1382, localized in the outer membrane, is predicted to play a role in the Type II and Type III secretion systems, while PA3512 has a putative role in ABC transporter systems.

**Fig 6.**
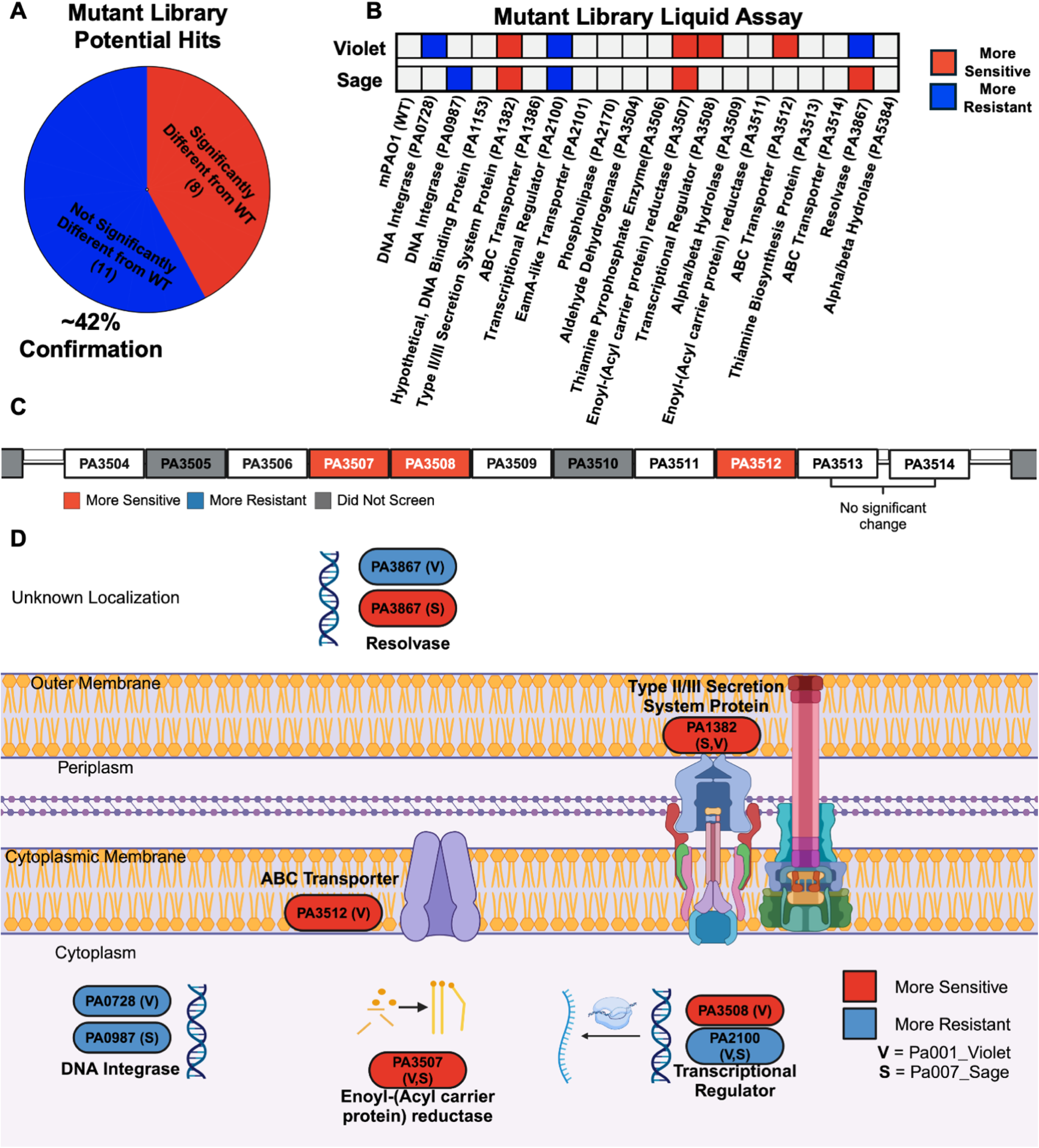
*P. aeruginosa* genes that affect phage infection. A) Forty-two percent (8/19) confirmed with findings from the bi-directional BLAST. B) Heatmap summary of mutant library liquid assay screen. A total of 19 different genes were tested. C) Genome region containing PA3504 – PA3514 in *P. aeruginosa* PAO1. Bi-directional BLAST revealed several genes from this localization that could affect phage infection. Three genes (3507, 3508, and 3512) significantly increased sensitivity to phage. D) Diagram of predicted location and functional annotation of the putative hit genes involved in phage infection. Most genes (5/8) encode proteins that remain in the cytoplasm but interact with the outer membrane or metabolic activities. One gene, PA1382, is found in the outer membrane. Red indicates mutants that were more sensitive to the phage; blue is resistant. “V” denotes Pa001_Violet, while “S” denotes Pa007_Sage. Gene annotations were obtained from BV-BRC.

To further assess whether introducing random mutations is enough to alter bacterial susceptibility to phages, we subjected two *P. aeruginosa* strains, mPAO1 and CDC 770, to UV-induced mutagenesis. While mPAO1 is sensitive to Pa001_Violet, the CDC 770 strain is resistant to this most robust phage. We tested phage specificity against ten randomly mutagenized isolates of each bacterial strain using the liquid assay and compared the results to the non-mutagenized parental strain. We found that four out of 20 mutagenized mPAO1 isolates showed a significant increase in phage sensitivity, while no changes were observed for CDC 770 (S14 Table). These results indicate that simply introducing random mutations does not affect bacterial susceptibility to phages, suggesting that the eight bacterial components identified by bi-directional BLASTp analysis play a specific role in regulating phage infection. Collectively, we identified genes that are potentially involved in phage infection and could offer new opportunities to improve or predict the efficacy of phage therapy. Our approach can be employed to identify genes across other species.

## Discussion

While municipal wastewater is a rich reservoir for phages and their bacterial hosts (37,38), tapping into hospital wastewater allows one to search for phages that may target bacteria that are endemic to the local area. The source of our phages may also explain why they were effective at targeting bacteria from the same hospital. Given that hospitals are reservoirs of more resistant variants of *P. aeruginosa*, our results suggest that phages may prefer to infect such strains, possibly because of increased expression of efflux pumps or decreased fitness. Such an isolation approach and subsequent application could potentially help control nosocomial *Pseudomonas* outbreaks, which often become widespread and cost hospitals millions of dollars (7,39–41). Our iterative screening approach successfully yielded phages capable of targeting a broad spectrum of *P. aeruginosa* from diverse sources. Surprisingly, however, the standard three rounds of phage isolation commonly reported in the literature did not result in a homogenous isolate (42,43). These results highlight the need for more stringent or extended purification steps to ensure clonal phage populations, especially when considering their therapeutic or experimental applications.

Genome sequencing and TEM imaging identified diverse phage genera classified into several distinct families and morphologies, including *Chimalliviridae Phikzvirus* (Pa001_Violet and Pa007_Sage), the genus *Bruynoghevirus* (Pa003_Juniper), *Skurskavirinae Pakpunavirus* (Pa003_Lilly), *Migulavirinae Litunavirus* (Pa004_Willow), *Jondennisvirinae Septimatrevirus* (Pa005_Rush), and *Peduoviridae Citexvirus* (Pa006_Rose). The identified phages displayed a variety of structural features. For example, the two *Phikzvirus* members, *Pakpunavirus*, and *Citexvirus* are characterized by icosahedral heads and contractile tails, two members of the *Litunavirus* and *Bruynoghevirus* genera are characterized by short, non-contractile tails and icosahedral heads, and a single member from the *Septimatrevirus* genus is characterized by a long non-contractile tail and icosahedral head. Notably, many of these genera are specific to *P. aeruginosa*. For example, phage phiKZ is known for its large genome and the ability to infect a broad range of *P. aeruginosa* strains (44). It is, therefore, not surprising that our two phiKZ-like phages, Pa001_Violet and Pa007_Sage, demonstrated the most robust performance, characterized by their strong lytic activity and efficacy against multiple strains, including those with multidrug resistance. Among our other identified phages, Pa003_Lilly (*Pakpunavirus*) (45), Pa006_Rose (*Citexvirus*) (46), Pa004_Willow (*Litunavirus*) (47), Pa002_Juniper (*Bruynoghevirus*) (48), and Pa005_Rush (*Septimatrevirus*) are all known to infect *P. aeruginosa* (49).

Out of the seven phages, two, Pa001_Violet and Pa007_Sage, were able to target over 50% of the CDC *P. aeruginosa* collection that primarily contains MDR and pandrug-resistant strains. These two giant phiKZ-like phages are known for their resistance to bacterial antiviral mechanisms through the formation of a DNA-shielding nucleus (50), which likely contributes to their exceptional potency in our *in vitro* and *in vivo* experiments (51). Despite the robust efficacy of phiKZ phages and unique resistance mechanisms (52), very limited studies exist using animal models, and none were published in humans despite being used in commercial products in Georgia and Russia (44).

Our stringent criteria used to characterize phages rely on AUC, RiMA, and TURU—three metrics that measure specificity and ensure a robust and comprehensive assessment of phage efficacy and bacterial susceptibility. Utilization of such a tripartite approach is essential as it captures different aspects of bacterial proliferation, metabolism, and response to phage treatment. For example, some bacterial strains may exhibit minimal changes in one metric (e.g., growth curve) but significant changes in another (e.g., metabolic activity). Therefore, assessing several measurements simultaneously eliminates potential bias, detects subtle differences, and ensures accuracy. Furthermore, our colorimetric approach records metabolic activity, providing a close assessment of proliferation but not direct bacterial viability. For example, we frequently notice that bacteria are killed in wells that completely reduce the MTT dye (S5 Fig), indicating that the signal develops before the bacteria are infected and killed. Hence, a small but significant change across our three metrics can result in robust killing.

Our results highlight a slight difference in phage efficacy in liquid versus solid media, a result that is likely affected by several contributing factors, including phage titer, MOI, nutrient availability, bacterial growth rates, phage dynamics, and biofilm formation. Given that an aqueous environment is more physiologically relevant to infections *in vivo*, our approach provides a promising tool for assessing phage efficacy, which is further supported by our mouse experiment. Despite the reduced concentration (1/7^th^) of each of the seven combined phages, their efficacy was better than each individual phage alone, as highlighted by the differences in RiMA scores between the cocktail and individual phages (Fig 3C). These results are consistent with *P. aeruginosa* phage cocktails published across several studies (53–55), emphasizing increased constraint on the evolution of phage-resistant bacteria. The enhanced efficacy of our cocktail could have also occurred from the iterative screening approach we took to isolate phages, which warranted the selection of diverse phages as demonstrated by their characterization and antimicrobial properties. The efficacy of phages revealed by the *in vitro* colorimetric assay employed for phage characterization translates into *in vivo* models. The experimentally derived cocktail successfully treated *P. aeruginosa* infection in mice, almost completely clearing bacteria from all organs and rescuing animals from lethal bacteremia. These results emphasize the reliability of our *in vitro* assessment in predicting *in vivo* efficacy.

*P. aeruginosa* mutants of five genes enhanced the sensitivity to two robust phages, Pa001_Violet and Pa007_Sage (Fig 6B). Interestingly, these genes included a Type II/III secretion system protein (PA1382), phage DNA invertase/resolvase (PA3867), an Enoyl-(Acyl carrier protein) reductase (PA3507), a transcriptional regulator (PA3508), and a predicted ABC transporter protein (PA3512). Interestingly, PA3867 was observed to increase sensitivity for Sage while decreasing sensitivity for Violet, highlighting potential differences in infection mechanisms between the two viruses. Genomic rearrangements in *Lactobacillus johnsonii*, such as the integration of novel prophages and the presence of CRISPR elements, have been implicated in strain-specific phage defense mechanisms (56). It is, therefore, possible that the identified DNA invertase/resolvase (PA3867) facilitates resistance to our phiKZ-like phage Violet through similar genomic rearrangements. These same rearrangements, however, increased sensitivity to Sage. Efflux pumps and other transmembrane transport systems often function as phage receptors (57); with the functional redundancy between membrane transporters, the inactivation of one system enhances the expression of another compensatory transport (58), which could serve as a phage receptor. Regarding PA1382 and its putative interactions in the Type II/III secretion system pathways, changes in phage infection efficacy highlight interesting differences between the mutants and wild type. Elimination of PA1382 increased sensitivity to both Violet and Sage. This change in infection could be related to a compensatory effect with the elimination of critical secretion pathway proteins. However, Type II and Type III secretion systems have been observed to have unique interactions with bacteriophages in other bacterial species. In *Vibrio cholerae,* a filamentous phage CTXΦ utilizes EpsD, the outer membrane component of the Type II secretion system, to escape the bacterium (59). Additionally, in *Salmonella enterica* serovar Typhimurium, a portion of the Type III secretion system is encoded by a prophage, increasing virulence (60). These interactions highlight potential superinfection exclusion that could occur during phage infection, as elimination of PA1382 in mPAO1 significantly increased phage infection sensitivity. A transcriptional regulator (PA3508) increased sensitivity to phage Violet. This regulator, which has been linked to biofilm formation and lipopolysaccharide (LPS) synthesis, most likely alters phage infection through structural changes (61,62). LPS is a known target for many phages (63–65); additionally, alterations in LPS structure are known to increase resistance to phage in various *Pseudomonas* species (66,67). PA3508 was originally identified in the phage-sensitive isolates due to low sequence similarity to PAO1. Thus, it is conceivable that these isolates have alterations in their LPS structure, thus increasing sensitivity to infection by Violet. Finally, PA3512, a putative ABC transporter, may have several interactions with phage infection. First, PA3512 may be linked to prophage modification of *P. aeruginosa* LPS. Using a strain of *Escherichia coli* K12 transformed with a plasmid containing O-antigen modification genes from *Raoultella terrigena*, it was shown that ABC transporters are required for successful glucosylation (68). ABC systems have also been linked to A-band LPS transport in *P. aeruginosa* (69). As discussed, modification of the LPS structure could alter phage infectivity, as LPS is a known target for phage infection. Second, ABC transporter systems have been linked to both the uptake and export of antibiotics (70,71). If the observed change in infection is related to ABC systems that govern antibiotic resistance, phage Violet would be an effective candidate for phage-antibiotic synergy applications.

The bi-directional BLASTp approach also yielded four genes whose mutants increase resistance to our phiKZ-like phages. Interestingly, a transcriptional regulator (PA2100) is located adjacent to the efflux transporter (PA2101) and is known for the regulation of several efflux pumps (72), including MexAB, which component was recently identified in a screen for *P. aeruginosa* mutants that confer phage resistance (73). Furthermore, the MexAB-oprM system has been previously described as *P. aeruginosa* phage receptor (57). In our liquid assay, the elimination of PA2100 greatly increased phage resistance to both Violet and Sage. As PA2100 is a negative regulator of the MexAB-oprM efflux pumps, the elimination of PA2100 is expected to increase their prevalence in the bacterium. Thus, these results further suggest that the MexAB-oprM efflux pump is involved with phage resistance as our phikz-like phages experienced reduced efficacy. As a transcriptional regulator, however, PA2100 could be involved with other functions within the cell that affect phage infectivity. Though the mutants became more resistant to Violet and Sage, it should be noted that this increase in resistance was only observed after 22 hours of incubation. At 12 hours, metabolic activity of both the wild type and mutant were still reduced by nearly 90%. Notably, this efflux pump is one of the mechanisms contributing to carbapenem resistance in *P. aeruginosa* (74). If an increase in these efflux pumps do not severely inhibit phage infection, potential phage-antibiotic synergy applications could occur with antibiotic treatment to increase the efflux pump density. A predicted DNA integrase (PA0987) was previously found among other MvaT and MvaU-regulated genes that facilitate induction of the Pf4 prophage (75). Therefore, loss of PA0987 can possibly stabilize the Pf4 lysogenic state and prevent infection by other phages, likely through superinfection exclusion mechanisms. Interestingly, the mutant of intF4 (PA0728), a phage DNA integrase (76), only affected susceptibility to Violet, while PA0987 affected Sage. Together, these results highlight the need for effective phage cocktails to reduce the risk of mutations inhibiting successful phage infection. When identifying potential genes that affect phage infection, we used seven cocktail-resistant isolates. As we only screened two of the seven phages from the cocktail, it is possible that these genes identified could have altered infection differently for the other phages. While further work is needed to decipher the role of the identified genes in phage infection, our bi-directional BLASTp approach combined with liquid assay can be employed for a rapid, high-content assessment of bacterial genes that affect phage infection.

Our study reveals a stringent and robust assessment of phage antimicrobial efficacy *in vitro* and a good predictor of their activity *in vivo*. A combination of phenotypic-genotypic analyses identified previously unexplored *P. aeruginosa* genes involved in phage infection. Despite the need for additional research to determine the precise function of these bacterial products, a comprehensive understanding of their role in phage infection is crucial for advancing phage therapy.

## Materials and Methods

### Bacterial strains

Unique *Pseudomonas* isolates with varying resistance patterns were identified by the clinical infectious diseases team at UFHealth Shands Hospital and communicated to the hospital’s clinical microbiology lab. The isolates were streaked on fresh TSA plates for each testing and subjected to DNA extraction for whole genome sequencing. We obtained a panel of 55 *P. aeruginosa* clinical isolates from the Centers for Disease Control and Prevention (CDC) and the Food and Drug Administration (FDA) Antimicrobial Resistance Isolate Bank. Environmental and animal bacterial isolates were previously isolated and published (77). A full list of bacterial strains used in this study can be found in S15 Table.

### Growth of bacterial stocks

All bacterial stocks used in this study were stored at −80 °C in 15% glycerol. For every experiment, fresh bacteria were inoculated into 5 mL of Lennox Broth (LB) and incubated overnight at 37 °C while shaking at 220 revolutions per minute (RPM).

### Isolation of bacteriophage

Untreated raw wastewater samples were collected from the water reclamation facility and from upstream of the UF water reclamation facility from a gravity main that collects wastewater from the UFHealth Shands Hospital North Tower. Samples were collected in 1 liter precleaned high-density polyethylene bottles by attaching the bottle to a collection pole with a clamp. The bottle was lowered into the waste stream and allowed to rinse three times before collecting the final sample. The samples were immediately placed in a cooler for transport to the lab, where they were stored at 4 °C until further analysis. Fifty milliliters of wastewater were centrifuged at 2,000 xg for 5 minutes to remove large particulate matter. Centrifuged wastewater was then filtered through a 0.22 µm membrane to remove unwanted bacterial contaminants. Filtered wastewater was mixed 1:1 with 35 mL of fresh 2x tryptic soy broth (TSB) and 1 mL of overnight bacterial host. The wastewater/bacterial mixture was incubated overnight at 37 °C while shaking at 220 RPM. After incubation, phage lysate was centrifuged at 10,000 xg for 20 minutes to pellet bacteria. Centrifuged phage lysate was filtered through a 0.22 µm syringe filter to remove any bacteria remaining in the supernatant. Filtered phage lysate was plated with the bacterial host in a double-layer agar assay (see procedure). Following incubation, a single plaque-forming unit (PFU) was picked using a pipette tip and transferred to a fresh double-layer agar plate with the bacterial host. This picking isolation technique was repeated until each phage had undergone nine rounds of purification. After the last round of isolation, a single plaque was picked and inoculated into 35 mL of 2x TSB containing 1 mL of overnight host bacteria and incubated overnight at 37 °C while shaking at 220 RPM. Following the incubation, phage lysate was filtered through a 0.22 µm vacuum filter and concentrated by centrifugation at 22,000 xg. The supernatant was aspirated, and the phage pellet was resuspended in 6 mL of phage buffer (10 mM Tris-HCl, pH 7.0; 10 mM MgCl_2_; 5 mM CaCl_2_). A 1 mL aliquot was frozen at −80 °C as a stock, and the remaining was stored at 4 °C for subsequent experiments. Each phage was initially screened against the AR Isolate Bank collection of 55 clinical strains of *P. aeruginosa*, and phage-resistant bacteria were used as hosts to isolate additional phages. This process was repeated until seven phages had been isolated, covering approximately 90% of the CDC library.

### Bacteriophage sequencing and analysis

Bacteriophages were concentrated by high-speed centrifugation (∼22,000 xg, 60 minutes, 4°C). To remove bacterial nucleic acids, concentrated phage stocks were treated with 0.001 U DNase I (ThermoFisher) and 2 µg/mL RNase A (NEB) and incubated at 37°C for 90 minutes, followed by inactivation for 10 minutes at 75 °C. Phage DNA was extracted using a Monarch® Genomic DNA Purification Kit (#T3010S) and sequenced using the Illumina platform at SeqCenter (PA, U.S.). The reads were analyzed using the FASTq suite and trimmed using Trim Galore on the Bacterial and Viral Bioinformatics Resource Center’s (BV-BRC) (78). Genome assembly was performed using SPAdes 3.15.5 and corrected using Pilon with default parameters (79,80). Phage genomes were compared to their closest identified relatives using BLAST, progressiveMAUVe, and Clinker (81–83). Annotation of phage genomes was completed using Prokka (local) and VIGOR4 (BV-BRC) (84,85). Identification of phage integrases and phage toxins was completed by utilizing the automated annotation output of Prokka within clustered BLASTp analyses.

### Bacterial genome sequencing and analysis

Bacterial DNA was extracted from each isolate using a Qiagen DNeasy Blood and Tissue Kit (Qiagen, Valencia, CA) according to the manufacturer’s instructions. The DNA libraries were constructed using the Nextera XT sample preparation kit (Illumina, San Diego, CA) following manufacturer’s protocol. Whole genome sequencing was conducted using Illumina MiSeq 250 bp paired-end sequencing. The reads were analyzed using BV-BRC (78). The quality of reads was evaluated using FastQC and trimmed using Trim Galore (86). Raw Illumina reads were assembled using SPAdes and annotated with RAST took kit (RASTtk) (87,88). All analyses were run with default parameters. The whole genome sequences have been deposited to Sequence Read Archive (SRA) and the corresponding Accession Numbers are available in S15 Table.

### Identification of bacterial isolates in collection

The whole genome sequence FASTA files were analyzed using Barrnap (Bacterial Ribosomal RNA Predictor) to identify and extract predicted 16S rRNA gene sequences. Barrnap was run with default parameters. The extracted 16S regions were subsequently submitted to BLASTn (Basic Local Alignment Search Tool nucleotide) against the NCBI 16S ribosomal RNA database for taxonomic identification at the genus and species levels. The BLASTn results were analyzed to determine the closest taxonomic matches based on sequence similarity scores.

### High-content liquid assay screen

The *in vitro* liquid assay screen was performed using OD_600_ and PFU-normalized concentrations of bacteria and phages, respectively. A working solution of 500 µg/mL MTT was prepared in fresh LB medium supplemented with 5 mM CaCl_2_ and 10 mM MgCl_2_. Overnight bacterial cultures were diluted to a concentration of 2×10^4^ CFU/mL in the MTT working stock. Subsequently, phages or phage cocktails were added at the concentration of 1×10^4^ PFUs per mL (PFU/mL), achieving an initial multiplicity of infection (MOI) of 0.5. The bacterial and phage mixtures were then aliquoted into a 96-well plate, with each well containing a total of 100 µL, resulting in eight technical replicates per treatment group per phage. Each experiment was repeated twice for a total of 16 replicates. The 96-well plate was incubated for 22 hours at 37 °C in the OmniLog® system and the phage efficacy was evaluated based on the relative unit output. Phages that exhibited at least a 10% reduction in RiMA (approximately 3x standard error of the control), a significant difference in the AUC based on the Student’s *t*-test, and a significant reduction across at least two relative unit measurements for each treatment group were deemed effective.

### Double-layer agar assay

The assay was performed to assess the lytic activity of the isolated phages against selected bacterial strains. Briefly, a soft agar overlay was prepared by diluting 1 mL of phage lysate to the desired dilutions. One hundred microliters of the diluted phage were mixed with 1 mL of an overnight bacterial culture, which was normalized to an optical density at 600 nm (OD_600_) of 0.1. The phage-bacteria mixture was then combined with 2 mL of 1% molten agar and immediately poured onto a fresh TSA plate to form a 0.7% top agar layer. After the top layer solidified, the plates were incubated at 37 °C for approximately 16 hours to allow for plaque formation.

### Solid assay phage spotting

Bacterial hosts used in solid assay experiments were grown for 12-16 hours at 37 °C with shaking at 220 RPM. The cultures were standardized to OD_600_ of 0.1, mixed with 1% molten agar, and poured over a fresh TSA plate to form approximately 0.7% top agar layer. Ten microliters of each phage and phage cocktail, at a concentration of 1×10^7^ PFU/mL, were spotted in their designated section on the bacterial lawn. The plates were incubated overnight at 37 °C and examined for plaque formation. Plaque development was classified into three categories: Sensitive (clear plaques), Intermediate (slight inhibition with some bacterial survival within the plaque), and Resistant (no plaque formation).

### Transmission electron microscopy

Imaging of all phages was acquired using an FEI Spirit 120kV transmission electron microscope. Formvar carbon film grids with copper mesh were cleaned using glow discharge in a PELCO easiGlow Discharge Cleaning System. Bacteriophage stocks were concentrated using high-speed centrifugation (∼22,000 xg, 1 hour, 4°C), and 10 µL of phage stocks were spotted on the carbon grids. After incubation at room temperature for ∼40 seconds, one drop of 0.5% uranyl acetate was dripped from a 0.22 µm syringe filter onto the grid and incubated while covered for ∼30 seconds. Grids were gently tapped against filter paper to remove excess liquids, covered, and placed in a dark, ventilated location to dry overnight for imaging the next day.

### UV mutagenesis

Two strains of *P. aeruginosa,* mPAO1 and CDC 770, were selected based on their susceptibility to Pa001_Violet phage – mPAO1 is susceptible, and CDC 770 is resistant. Host strains were inoculated from glycerol stocks and grown for approximately 16 hours at 37 °C with shaking at 220 RPM. The cultures were then diluted to OD_600_ of 0.1 and spread across fresh TSA plates using wood-handle cotton swab applicators. A Stratalinker was used to expose each plate to 3 seconds of UV radiation, 1 mW/cm^2^ (3 mJ/cm^2^). This exposure to UV radiation reduced bacterial lawn to individual colonies for both mPAO1 and CDC 770. Ten bacterial colonies from each original strain were selected and screened against Pa001_Violet phage using the liquid assay.

### Mouse bacteremia model

Ten female C57BL/6 mice, aged 8-12 weeks, were divided into two groups: “no phage” and “phage,” with five mice in each group. All mice received an intraperitoneal (IP) injection of 1×10^6^ CFU *P. aeruginosa* PAO1. Thirty minutes post-infection, mice in the phage group were IP-inoculated with a phage cocktail containing a total of 1×10^9^ PFU in phage buffer. The phage cocktails were diluted in phage buffer (10 mM Tris HCl, 10 mM MgCl_2_, 5 mM CaCl_2_, pH 7). Control mice were treated with sterile phage buffer alone. Treatments were readministered every 12 hours post-infection. At 36 hours post-infection, the mice were sacrificed, and their organs (kidneys, spleen, liver, lungs, heart) were harvested, weighed, homogenized, and plated for CFU quantification on Pseudomonas isolation agar (Millipore #17208). Before dissection, the abdominal cavity was washed with 10 mL of PBS, which was collected for subsequent dilution and CFU plating. Each harvested organ was placed in a whirl-pak and resuspended in 10 mL PBS supplemented with 1% Triton-X, followed by homogenization. The homogenates were serially diluted six times at a 1:10 ratio in phosphate-buffered saline (PBS). Ten microliters of each dilution were plated on Pseudomonas isolation agar and incubated overnight at 37 °C. Following incubation, colonies were counted to determine the average CFU/gram of tissue. Each organ dilution was plated in six replicates. With five mice per group and six replicates per organ, a total of 30 replicates per organ were analyzed.

### Phylogenetic analysis

The phylogenetic tree was constructed using the Bacterial Genome Tree tool on BV-BRC, based on an alignment of 500 single-copy genes. The tree was visualized with iTOL (89), and bacterial strains resistant to the phage cocktail were highlighted in blue.

### Bi-directional BLAST

Following the identification of bacterial genomes of interest through the phylogenetic tree, a proteome-wide bi-directional BLASTp was conducted using the Proteome Comparison tool on BV-BRC. This analysis included a clade of seven cocktail-resistant *P. aeruginosa* isolates (CDC 230, 242, 243, 248, 249, 254, and 255), two sensitive isolates (CDC 264 and K016), and *P. aeruginosa* PAO1 as the reference. The sensitive isolates were selected due to their close phylogenetic proximity to the resistant isolates. The BLASTp analysis CSV output of all annotated proteins corresponding to the selected bacterial genomes was used to arrange proteins by their similarity to PAO1, and those with 50% or lower scores were deemed as potentially involved in phage infection. The proteins selected as putative hits had to have a sequence similarity below the threshold consistent between the entire cluster of resistant and the selected sensitive strains. To confirm the predicted hits from the bi-directional BLASTp, we employed a transposon knockout library (90), and tested each of the 19 identified genes with low sequence similarity score (<50%) in a high-content liquid assay using Pa001_Violet and Pa007_Sage phages and employing the three liquid assay metrics – RiMA, AUC, and TURU. Mutants that exhibited significant differences in phage infection compared to wild-type were deemed as genes involved in phage infection.

### Mutant Library Screen

Predicted mutants from the bi-directional BLAST were selected from the Manoil Transposon Knockout Library and grown for approximately 16 hours at 37°C while shaking at 220 rpm in 5 mL of fresh LB inoculated with 5 µg/mL tetracycline, as recommended by the Manoil Library guidelines. After incubation, mutants were standardized to OD_600_ = 0.0001 in 5 mL of MTT/LB. Pa001_Violet and Pa001_Sage were diluted and added to bacteria for an initial MOI of 0.5 and both no-phage and phage treatments were added to a 96 well plate using the same methods as the liquid assay. After incubation in the OmniLog, the mutant metabolic curves were compared to mPAO1 as control using the three metrics: RiMA, AUC, and TURU. Mutants that exhibited significant difference in all three metrics across at least two biological replicates (16 technical replicates) were considered significant.

### Statistical analyses

Graphs were generated using GraphPad Prism 10 (v10.2.4) for Mac, GraphPad Software, Boston, Massachusetts USDA, www.graphpad.com. Statistical significance was determined using unpaired Student’s *t*-test, Multiple *t*-tests, or Welch’s *t*-test, depending on the data set. Error bars represent the standard error of the mean. Significance levels were denoted as follows: *, p<0.05; **, p<0.01; ***, p<0.001, and ****, p<0.001.

## Acknowledgments

We thank Ms. Anthea Sabol for isolates from the Shands microbiology lab. Special thanks go to the CDC and FDA’s Antibiotic Resistance (AR) Isolate Bank for generously providing *P. aeruginosa* clinical isolates used in this study. Figures 1, 4, and 6 were created using a paid version of biorender.com. We thank the University of Florida Interdisciplinary Center for Biotechnology Research (ICBR) Electron Microscopy core (RRID:SCR_019146) for providing services to image phages.

## Ethics declarations

The authors declare no conflicts of interest. All the animal work was performed under prior approval from the University of Florida Institutional Animal Care and Use Committee (IACUC number 202200000705). The *P. aeruginosa* isolation protocol from patients was reviewed and approved by the University of Florida Institutional Review Board (IRB202000185). De-identified bacterial isolates were collected from patients and no consent was obtained.

## Supporting Information

**S1 Table. *P. aeruginosa* CDC isolate characterization.** The antibiotic classification for sensitive, intermediate, or resistant as well as the antibiotic resistance mechanisms, were provided by the CDC’s AR Isolate Bank (Panel ID 12). Intact prophage regions were recorded as described by PHASTER. Strains that are highlighted blue are resistant to all phages used in this study. Amikacin (AN), Aztreonam (ATM), Cefepime (FEP), Ceftazidime (CAZ), Ceftazidime/avibactam (CAZ-AVI), Ceftolozane/tazobactam (C/T), Ciprofloxacin (CIP), Colistin (CL), Doripenem (DOR), Imipenem (IPM), Levofloxacin (LVX), Meropenem (MEM), Piperacillin/tazobactam (TZP), Tobramycin (NN).

**S2 Figure. Clinker alignments of phage genomes.** The alignments for Violet (A), Willow (B), Rush (C), Rose (D), and Sage (E) are given as compared to their closest identified relatives. The alignments for Juniper and Lilly are shown in **Figure 1**. Black regions between the aligned genes represent high identity (%), while grey is representative of lower identity (%).

**S3 Movie. Liquid assay MTT reduction video.** A time-lapse video of a sample plate from the OmniLog® screens showing the difference in reduction between no-phage and phage-treated wells. The top portion of the plate (Rows A-D) is not treated with phage, while the bottom half (E-H) is treated. Each column represents a unique bacterial isolate. Darker wells indicate reduction (metabolic activity) and result in higher relative unit outputs. The video was annotated using Camtasia (Version 2023.3.14)

**S4 Fig. OmniLog® error RiMA cutoff.** Reduction of metabolic activity of a control (no phage) *P. aeruginosa* PAO1 from 164 wells across 22 separate plates. The average standard error of the mean is 3.20. The RiMA cutoff of ten is approximately three times the average standard error of the control.

**S5 Fig. Elimination of bacteria from reduced wells.** *P. aeruginosa* PAO1 was grown in various dilutions ranging from MOI 5,000 to MOI .0005 to visualize well reduction in the presence of phage. A) The metabolic curves for plate wells A01, A05, and A06, representing bacterial control, MOI 5, and MOI 0.5, respectively for Sage infection in the liquid assay are shown. The metabolic activity (reduction) of the wells increases at 22 hours. B) Omnilog® plate image from the experiment at 22 hours. Wells A01, A05, and A06 appear reduced. C) Contents from wells A01, A05, and A06 were plated in a CFU assay. Plating revealed elimination of bacterial cells, as compared to control, for MOI 5 and 0.5, highlighting phage killing potential even despite the presence of tetrazolium dye reduction at 22 hours.

**S6 Fig. *P. aeruginosa* susceptibility to phage cocktail in solid and liquid assays.** The sensitivity of *P. aeruginosa* strains to phages was compared across two media types: solid and liquid. A) A phage cocktail, concentrated to 1×10^6^ PFU/mL, was spotted on a double-layer agar plate containing different isolates from the CDC library. Phage sensitivity, intermediate sensitivity, or resistance was recorded according to plaque formation and clarity on a plate. B) Results from the solid spotting assay and the colorimetric MTT assay were compared. Strains that did not match across both assays are highlighted in red. (n=2).

**S7 Table. Liquid assay of non-*Pseudomonas aeruginosa* isolates.** Analysis of the 47 non-*P. aeruginosa* isolates revealed that the cocktail could not inhibit any bacteria based on the three liquid assay metrics used. R – resistant, n – non-significant.

**S8 Fig. Phage cocktail specificity to *P. aeruginosa*.** A phylogenetic tree using 188 single-copy genes across 166 good-quality bacterial isolates was constructed using software on BV-BRC’s website (A). Bacterial isolates resistant to the cocktail are highlighted in blue. On the left side of the tree, all isolates in blue represent the non- *Pseudomonas aeruginosa* isolates. B) Sample metabolic curves and AUC analyses for two isolates, *Pseudomonas putida* and *Escherichia coli* are shown with their resistance to cocktail treatment.

**S9 Fig. Single phage-treated mouse vs control group.** The single phage-treated mouse that displayed *P. aeruginosa* colonies post-sacrifice was compared to the control group to examine bacterial reduction due to phage treatment. Across all organs, there was a significant reduction in bacterial colonies; the intraperitoneal (IP) wash had a single bacterial colony recovered. (A) The bacterial load (CFU/g tissue or CFU/mL IP wash) is listed for each organ, as well as the percent reduction between the control.

**S10 Movie. Mouse bacteremia model video.** Immediately prior to sacrificing, a video of both treatment groups (Control and Phage) was recorded. Compared to control mice, phage cocktail-treated mice exhibited no clinical signs of infection.

**S11 Fig. Predicted hits protein alignment.** To verify BV-BRC’s auto-annotation accuracy and assignment in the bi-directional BLASTp, alignments of all liquid assay-confirmed proteins were performed on BV-BRC using Mafft. All 8 hits—PA00728 (A), PA0987 (B), PA1382 (C), PA2100 (D), PA3507 (E), PA3508 (F), PA3512 (G), and PA3867 (H) from both the seven phage-resistant (CDC 230, 242, 243, 248, 249, 254, and 255) and two phage-sensitive (CDC 264 and K016) isolates, as well as PAO1 as reference are shown. All representative images are taken starting at residue 147.

**S12 Table. Predicted genes that influence phage infection.** Transposon mutants of each predicted gene were screened for phage sensitivity using Pa001_Violet and Pa007_Sage at an MOI of 0.5 in the liquid assay. To confirm increased sensitivity or resistance to phage per knockout, the three metrics were employed between phage treatment of the wild-type mPAO1 and the mutant (ΔRiMA, AUC, and TURU). To be considered significant, the mutant must have shown greater than 10 or less than −10 in ΔRiMA (highlighted light orange or light blue) and significance across AUC and TURU. RiMA and ΔRiMA scores are shown at 22 hours. Statistical significance was performed using the Student’s *t*-test (*, p<0.5; **, p<0.1; ***, p<0.001; ****, p<0.0001; ns, not significant). Δ - significant across at least two TURU thresholds. R – phage resistant (highlighted blue), S – phage sensitive (highlighted red).

**S13 Table. Predicted subcellular localization of the identified bacterial gene products that affect phage infection.** Using PSORTb version 3.03, the predicted localization and scores for each predicted gene product are given.

**S14 Table. Random mutagenesis of mPAO1 and CDC 770.** Freshly-swabbed plates of mPAO1 (phage sensitive) and CDC 770 (phage resistant) were exposed to UV and 10 mutagenized colonies (20 colonies total) were screened using Pa001_Violet in the liquid assay. To be considered significant, mutants must have significance across all three metrics. For RiMA in particular, the phage treatment of the wild-type was compared to the phage treatment of the mutant (ΔRiMA). Isolates must show scores greater than 10 or less than −10 to be considered more sensitive or resistant. Statistical significance was performed using the Student’s *t*-test (*, p<0.5; **, p<0.1; ***, p<0.001; ****, p<0.0001; ns, not significant). Δ - significant across at least two TURU thresholds. S – phage sensitive (highlighted red).

**S15 Table. Bacterial isolates used in the study.** Identification of sequenced isolates was performed by BLAST analysis of the 16S rRNA gene, as recovered by Barrnap, or the *rimM* gene, or similar gene if no *rimM* was annotated. The highest percent coverage and identity of each bacterium used are listed alongside the BLAST results. Blue rows with bold text indicate strains identified as non-*Pseudomonas aeruginosa* isolates. Orange rows with bold text indicate strains where no identification was obtained due to poor sequence quality. NS – not sequenced. *Isolate was identified by a clinical laboratory. Total of 184 bacterial isolates: 137 *Pseudomonas aeruginosa* isolates; 47 non-*aeruginosa* isolates.

